# Patients’ Derived Short-Term Glioma Culture: Identification of Aggressive, Drug-Resistant Phenotype

**DOI:** 10.1101/2020.06.04.131680

**Authors:** Syed Sultan Beevi, Vinod Kumar Verma, Manas Kumar Panigrahi, Aishwarya Bhale, Sailaja Madigubba, Radhika Chowdary, Radhika Korabathina, Sukrutha Gopal Reddy

## Abstract

**Background:** Clinical management of glioma is crucial irrespective of tumor grade. Despite newer treatment modalities, the prognosis of glioma is abysmal and, survival statistics are not remarkable. *In vitro* glioma culture is emerging as a standard model to get insight into phenotypic transformation, drug response, and tumor relapse. In this viewpoint, this study established comprehensive patient-specific short-term cultures comprising low-grade, and high-grade glioma, and evaluated their pertinence in the potential disease management.

**Methods:** 50 patients with MRI diagnosed glioma were recruited for this study. Primary glioma cultures established from fresh surgical tumor tissues, which were then evaluated for their intrinsic growth kinetics, response to temozolomide, and expression profile of Glial-Mesenchymal Transition (GMT) markers along with an oncogenic marker, cMyc.

**Results:** Short-term glioma culture was successfully established in 40 clinical samples. Glioma culture, irrespective of tumor grade, displayed two distinct patterns of growth kinetics – one with shorter doubling time (high-proliferating) and another group with longer doubling time (low-proliferating). Significant distinctive features were noticed between these two groups in terms of response to temozolomide, the expression pattern of GMT markers and their association with 1p/19q co-deletion and p53 expression.

**Conclusion:** Our findings effectively demonstrated the practicality of the development of short-term glioma culture toward a functional approach for personalized medicine. Our study revealed the presence of a highly proliferative, drug-resistant phenotype irrespective of tumor grade. Hence, short-term culture could be an important prognostic tool for predicting patient clinical responses and cue about imminent tumor relapse.

## Introduction

Malignant gliomas are the most common intra-axial primary tumors originating from the central nervous system’s supporting glial cells. Gliomas have the third-highest cancer-related mortality and morbidity rates among individuals worldwide (1) with an annual incidence of 1 million cases in India (2). The World Health Organization (WHO) classification system categorizes gliomas as grade 1 (biologically benign), grade II (diffuse astrocytoma/oligodendroglioma), grade III (anaplastic astrocytoma) and grade 4 (glioblastoma), based upon histopathologic characteristics such as cytological atypia, anaplasia, mitotic activity, microvascular proliferation, and necrosis (3).

The infiltrative phenotype has long been considered a contributory factor for clinical aggravation of glioma, which manifests as tumor recurrence and therapeutic intractability (4). Such infiltrative features have been explicitly linked to the phenomenon of epithelial-mesenchymal transition (EMT). The role of EMT in non-epithelial tumors like glioma is nonetheless uncertain. However, a recent classification of a mesenchymal subgroup of glioblastoma strengthens the notion that glial-mesenchymal transition (GMT) similar to EMT-like process may contribute to tumor progression, chemo-resistance, and tumor relapse regardless of tumor grade (5). Several studies have investigated the expression of EMT markers such as Vimentin, TWIST, SNAIL, E-cadherin, and N-cadherin in malignant gliomas and demonstrated their apparent role in invasion, drug resistance and tumor recurrence (6–9).

Patient-derived glioma cultures are gradually gaining impetus in brain tumor research and exemplify a way to get insight into prognostic and therapeutic attributes. Such *in vitro* models intently simulate the patient tumor’s biological traits, thus providing an opportunity to apprehend drug resistance, tumor recurrence, and response to therapy in a patient-distinctive manner (10).

Most of the *in vitro* glioma models referred to in the literature are exclusive of high-grade (11, 12) with little or no reference to low-grade glioma. However, low-grade gliomas are also equally fatal, even though they have better survival than patients with high-grade (13). Despite advancements in disease management, the overall prognosis of glioma per se is poor, impelling the need to devise an upfront approach for its comprehensive clinical management.

With this notion, we presumed that the development of comprehensive short-term glioma culture, including both low-grade and high grade and evaluation of its intrinsic growth kinetics together with GMT markers and drug response, could provide a simplistic approach for treatment and management of patients.

## Materials and methods

### Patients selection

Fifty patients with MRI diagnosed glioma, who visited the Department of Neurosurgery, Krishna Institute of Medical Sciences, Telangana, India, between December 2018 and January 2020, were recruited before their scheduled surgery. This study was approved by the Institutional Research Advisory Board and Ethics Committee. Written informed consent was obtained from all individual participants involved in the study. The study was conducted by following accepted guidelines for human material use and the 1964 Helsinki Declaration and its later amendments.

### Intraoperative tissue collection

Resection specimen of glioma tumor was divided into two parts, with one part in a sterile collection medium for the establishment of primary culture and other parts placed in buffered formalin for histological and pathological analysis.

### Establishment of primary glioma culture

Tumor tissues received in a sterile carrier medium was processed immediately within 2h upon arrival in the laboratory. Briefly, tumor tissue was minced by scalpel in PBS and digested with 0.1% collagenase I for 30 minutes at 37°C with intermittent shaking. Collagenase I activity was arrested by the addition of FBS and passed through a cell strainer (100 μm) to obtain a single-cell suspension. Cells were washed with PBS and seeded in T25 culture flask containing DMEM/F12 cell culture media supplemented with 10% FBS, 10ng/mL EGF/FGF-2, 2mM L-glutamine and 1% penicillin-streptomycin and allowed to adhere to their surface for 72 h at 37°C in 5% CO2. Unattached cells were removed through media change after 72 h, and the adherent cells were allowed to reach the onfluence of about 80% before being exposed to trypsin treatment and passaging. Cells were passaged 4-6 times and expanded for subsequent analysis.

### Morphological characterization of primary glioma cells

Glioma cells were cultured in T25 flasks to a confluence of 60 – 80% and morphology were assessed under an inverted phase-contrast microscope. Cells were photographed and processed using the cellSens software (Olympus, Japan).

### Population Doubling Time (PDT) Analysis

Cells were seeded at the density of 2 × 10^3^ cells per cm2 of surface area and incubated overnight at 37°C in 5% CO2. Plates were counted in duplicates every 24 h for seven consecutive days. Results were plotted on a linear scale as cell number versus time. Population doubling time (PDT) was calculated from the linear part of the curve using this equation: (t2 – t1)/3.32 × (log n2 – log n1), where t is time, and n is the number of cells.

### Sensitivity to Temozolomide (TMZ)

Cells (5 × 10^3^) were plated in 100μl DMEM/F12 media per well in triplicate in 96 well flat-bottom culture plates and allowed to adhere for 18h. Cells were then exposed to Temozolomide (in the concentration range of 2.5mM - 156μM) for 48h. An equal volume of DMSO was added to cells serving as a control. After 48h, cells were treated with 20μl of MTT (3-(4,5-dimethyl thiazolyl-2)-2,5-diphenyltetrazolium bromide) at a concentration of 5mg/mL in PBS and incubated for 4h. MTT containing media were discarded, added with 100μl of DMSO, vibrated for 10 min to dissolve the formazan crystals, and the absorbance was measured at 570nm by an ELISA microplate reader. The following formula calculated the degree of inhibition of cell proliferation in each sample: Inhibition of cell proliferation (%) = (1 – absorbance of the experimental group/absorbance of the control group) × Half maximal inhibitory concentration (IC50) was calculated by using regression analysis from GraphPad Prism.

### Expression pattern of GMT markers by Immunofluorescence

Glioma cells were cultured on 22×22mm square coverslips into 6-well culture plates and allowed to adhere overnight at 37°C in 5% CO2. Cells were washed in PBS, fixed for 20 min at room temperature in 4% paraformaldehyde in PBS. After fixation, cells were permeabilized with 0.1% Triton X-100 for 10 min and blocked with 5% bovine serum albumin for 1h at room temperature. Primary antibodies GFAP (rat anti-human IgM at 1:200 dilution), Vimentin (mouse anti-human IgM at 1:100), TWIST (mouse anti-human IgM at 1:100), E-cadherin (mouse anti-human IgM at 1:100), N-cadherin (mouse anti-human IgM at 1:100) and cMyc (mouse anti-human IgM at 1:100) were added and incubated overnight at 4°C. Cells were rinsed 3x with PBS and incubated with fluorescence-labeled (Alexa488/Alexa 594) secondary antibody and incubated at room temperature, protected from light 1h. Cells were rinsed 3x with PBS and mounted with VECTASHIELD mounting medium containing DAPI. The slides were analyzed by fluorescence microscopy (Olympus, Japan), and image processing was carried out using cellSens software. Cells were observed at different microscopic magnifications to make the precise assessment of percentages of stained cells. The entire slides were evaluated to avoid a biased selection of more or less intense staining areas. Quantitative scoring was done based on the percentage of positive cells and defined as: 0 for <10% positive cells (negative), 1 for 10 – 25% (low expression), 2 for 25 – 50% (moderate expression), 3 for 50 – 75% (strong expression) and 4 for >75% (intense expression)

### Pathological examination and immunohistochemistry of the patient glioma tissue

The tumor tissue was fixed in 10% buffered formalin, dehydrated with ethanol gradient, permeabilized with xylene, and paraffin-embedded. Then, serial 5μm thick sections were cut, deparaffinized with xylene, hydrated with ethanol gradient, and stained with hematoxylin and eosin (H & E). Meanwhile, immunohistochemical staining was performed for IDHR132, p53, and MIB-1 detection. Endogenous peroxidase inactivation was carried out with 3% H2O2, and antigen retrieval was performed in a microwave. Afterward, primary antibodies were added to each slide at appropriate dilutions, and the sections incubated with biotin-labeled secondary antibodies for 10 min. The final signals were developed using the 3,3’-diaminobenzidine substrate (DAB). The sections were analyzed by optical microscopy after counterstaining with hematoxylin.

### IDH mutational analysis

According to the manufacturer’s instructions, genetic DNA was isolated from FFPE tissue sections using QIAamp DNA FFPE tissue kit. The concentration of isolated DNA was determined with the anoDrop100. All the samples were analyzed for the mutation using the Qiagen IDH1/2 RGQ PCR kit, in the following loci: IDH1R132 (exon4), IDH1R100 (exon4) and IDH2R172 (exon 4). According to the manufacturer’s instruction, the desired genomic regions were amplified by qPCR using specific primers and probes.

### 1p/19q co-deletion

Co-deletion of 1p and 19q in tumor tissue samples were evaluated with fluorescence in situ hybridization (FISH) with locus-specific probes, LSI 1p36/1q25 or LSI 19q13/19p13. A positive result for 1p/19q co-deletion was assessed as the loss of 1p36 or 19q13 signal in more than 50% of nuclei.

### Statistical analysis

Clinical and biochemical data were expressed as mean ± standard deviation. Chi-square test for independence (χ2) and Pearson correlation coefficient (r) was applied to determine the association between proliferation status and clinical/pathological features of glioma. The *p*-value <0.05 was considered as statistically significant.

## Results

Fifty patients with MRI-diagnosed primary glioma were enrolled in this study. Out of 50 patients, five were excluded from the study as their later histological analysis revealed the presence of brain tumors other than glioma. Of the 45 patients, 14 were in grade II, 9, and 22 were in Grade III and IV, respectively. Detailed clinical and pathological characteristics of the glioma patients are presented as Supplementary Table 1.

### Primary glioma culture

Glioma tissue samples were digested enzymatically, and a single-cell suspension was grown as a monolayer culture in EFF/FGF-2 enriched media. The establishment of monolayer culture was successful in 89% (40/45) of the samples. Five samples (two in grade II & grade III and one sample in grade IV) either failed to attach or cease to grow after the first passage. Morphological evaluation, PDT, temozolomide sensitivity testing, and GMT markers analysis were performed mostly between passage 4 – 6 to minimize genomic instability due to the long-term culture process.

### Glioma culture showed mixed morphology

Primary glioma cells were micro-photographed to assess their morphology, and their representative images are depicted as Figure 1. Primary cultures showed cells with variable morphological features, predominantly having a dendritic-like and fibroblastic-like phenotype. There was no apparent difference in morphology between low-grade and high-grade culture.

**Figure 1.**
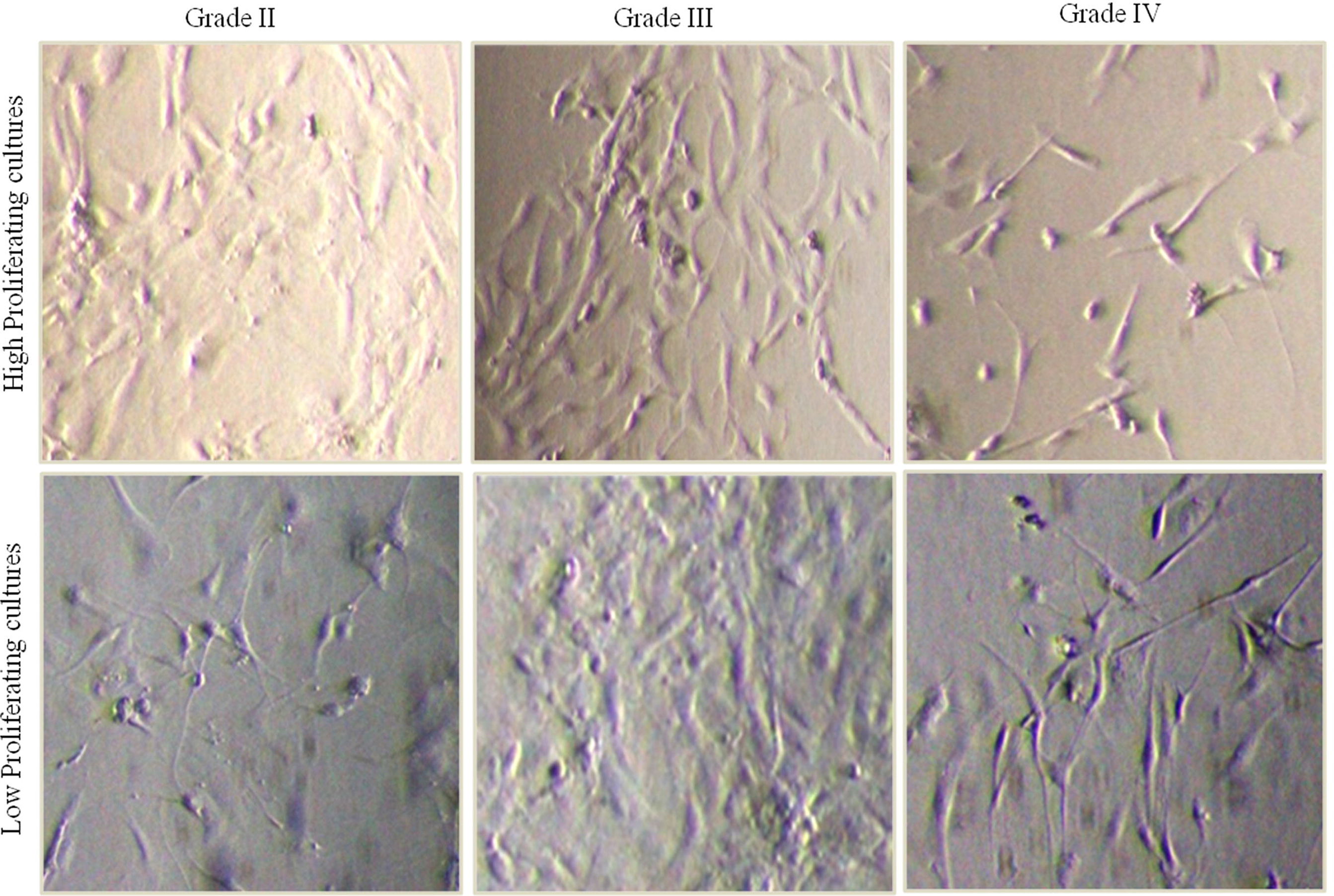
The phenotype of Primary Glioma Cells. Phase-contrast microscopic images depicting the morphology of glioma cells belonging to different grades

### Glioma culture displayed distinctive proliferation rate regardless of tumor grade

Estimation of Population doubling time (PDT) by the application of exponential growth equation (14) was employed to quantitate the proliferative capacity and intrinsic growth properties of the cultured glioma cells. Cultures were categorized as high-proliferating if PDT was less than 72h and low-proliferating if it was more than 72 h. Glioma cultures showed diverse proliferative capacity irrespective of tumor grades (Figure 2A). Seven out of 12 low-grade cultures exhibited high-proliferation, whereas this was observed in 12 out of 28 cultures in high-grade glioma. Furthermore, PDTs of high-proliferating high-grade gliomas were significantly shorter than those of low-grade gliomas, and it was contrasting in the case of low-proliferating cultures where high-grade gliomas exhibited longer PDTs as compared to low-grade glioma cultures (Figure 2B).

**Figure 2A.**
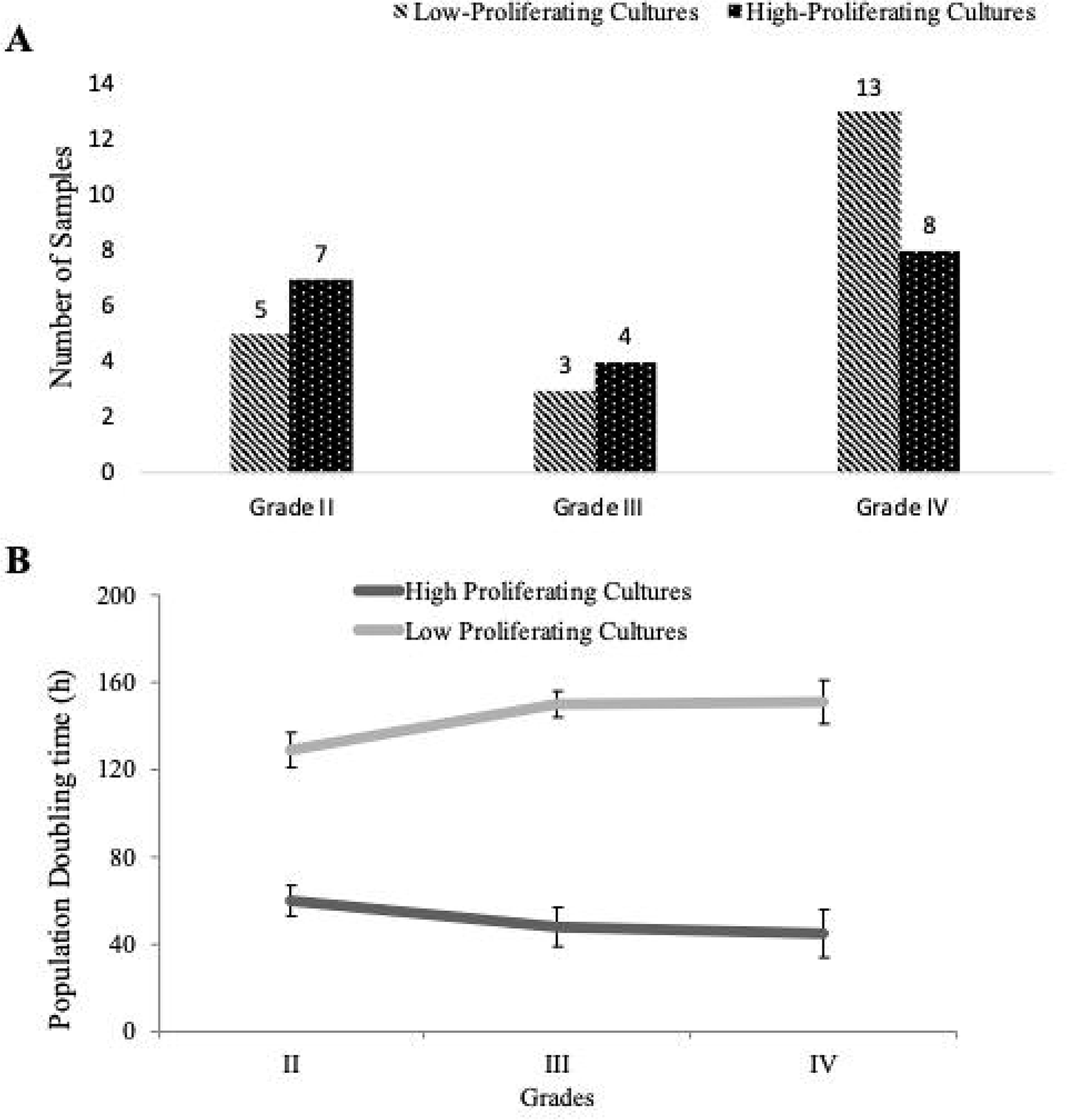
Grade-wise distribution of High and Low-Proliferating Cultures in absolute numbers. Cell cultures were assigned to the group with a high-proliferation rate when the doubling time was less than 72h, and the group had a low-proliferation rate when the doubling time was more than 72h.

**Figure 2B.** Population Doubling Time (PDT). Mean population doubling time is displayed grade-wise as a Line Graph. The doubling time for each sample was determined three times, and the values are represented as mean doubling time (h) ± standard deviation (SD). \**p* < 0.05; ** *p* < 0.01

### High-proliferating glioma culture was highly resistant to Temozolomide (TMZ)

The sensitivity of glioma cells was assessed towards temozolomide with its indicated concentrations. Inhibitory Concentration (IC50) value was calculated for all samples and categorized grade-wise concerning their proliferative capacity (summarized in Table 1). Samples belonging to the low-proliferating group were showing significantly lower IC50 as compared to high-proliferating cultures. Necessarily, IC50 of low-proliferating grade II and grade IV cultures were within the tested concentration of 0.156 – 2.5 mM compared with high-proliferating cultures where it was found to be in the range of 3 – 9 mM. Notwithstanding, all grade III cultures were highly resistant to TMZ with IC50 ranging between 4.5 – 13.3 mM, irrespective of their proliferation rate.

**Table 1.** Inhibitory Concentration (IC50) of Temozolomide in the Primary Glioma Cells by MTT assay.

| Primary Glioma Cells | IC50 of Temozolomide (mM) |  | *p* value |
| --- | --- | --- | --- |
|  | High Proliferating Cultures | Low Proliferating Cultures |  |
| Grade II | 3.110 ± 0.311 | 1.212 ± 0.209 | 0.00473 |
| Grade III | 13.29 ± 0.817 | 4.519 ± 0.532 | 0.00001 |
| Grade IV | 9.663 ± 1.134 | 1.447 ± 0.471 | 0.00001 |

Cell viability was measured after a 48h treatment with the indicated concentration of Temozolomide (as mentioned in the materials and methods. Results are the mean ± Standard Deviation (SD).

### High-proliferating glioma culture exhibited strong expression of Glial-Mesenchymal Transition (GMT) markers

Expression patterns of GFAP, vimentin, TWIST, E-cadherin, N-cadherin, and cMyc were evaluated in both high- and low-proliferating cultures and presented as Figure 3-8. High-proliferating cultures, irrespective of tumor grade, showed robust expression of GFAP, N-cadherin, and E-cadherin compared to low-proliferating cultures. Another exciting feature was the nuclear expression of cMyc in high-proliferating cultures instead of cytoplasmic expression in low-proliferating cultures.

**Figure 3:**
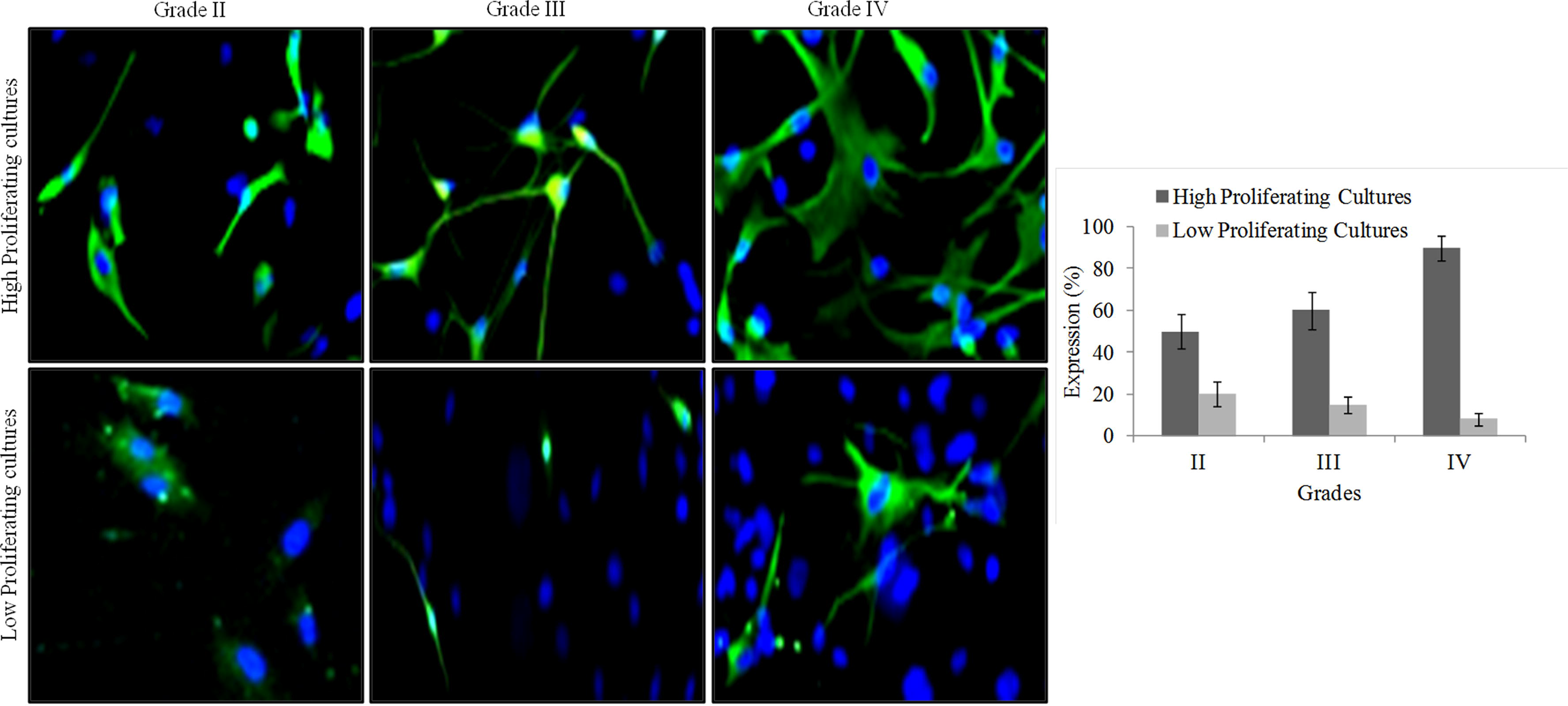
Representative Immuno-fluorescence staining of GFAP. Fluorescent microscope images show the cell nuclei labeled with DAPI (Blue) and GFAP labeled with respective antibody (Green, Alexaflour 488) and grouped grade-wise as per their proliferation rate. All images were captured at the Final Magnification of 100x.

**Figure 4:**
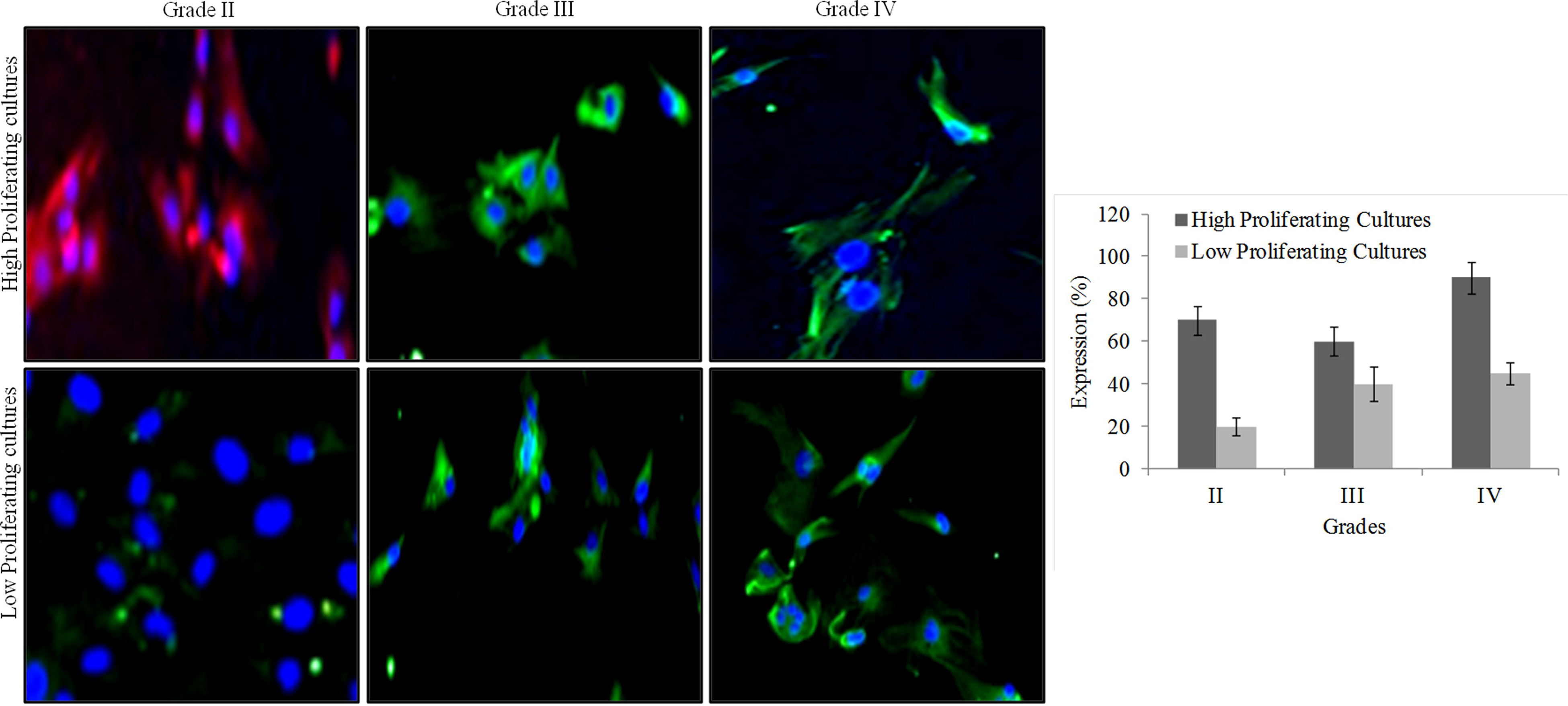
Representative Immuno-fluorescence staining of Vimentin. Fluorescent microscope images show the cell nuclei labeled with DAPI (Blue) and Vimentin labeled with respective antibody (Green, Alexaflour 488 and Red, Alexaflour 594) and grouped grade-wise as per their proliferation rate. All images were captured at the Final Magnification of 100x.

**Figure 5:**
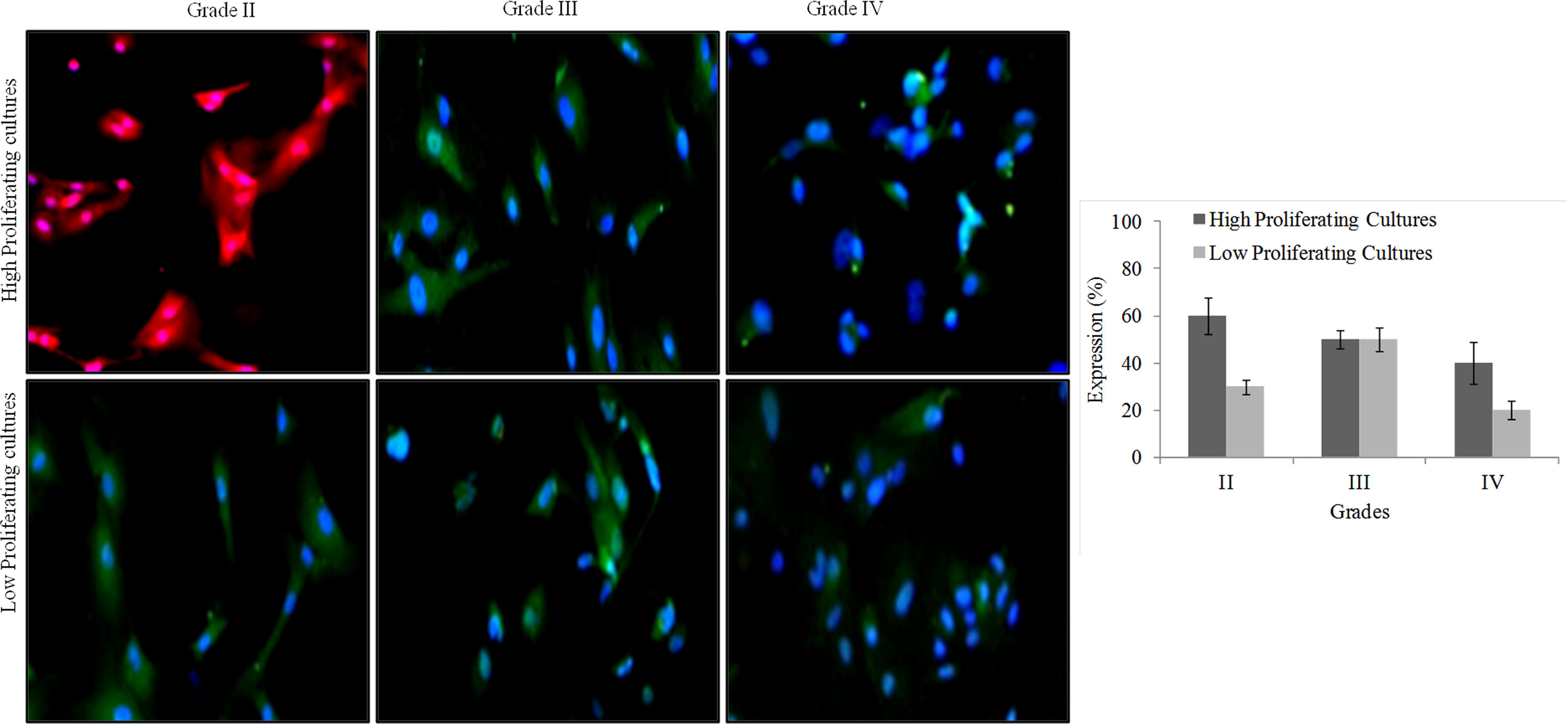
Representative Immuno-fluorescence staining of TWIST. Fluorescent microscope images show the cell nuclei labeled with DAPI (Blue) and TWIST labeled with respective antibody (Green, Alexaflour 488 and Red, Alexaflour 594) and grouped grade-wise as per their proliferation rate. All images were captured at the Final Magnification of 100x.

**Figure 6:**
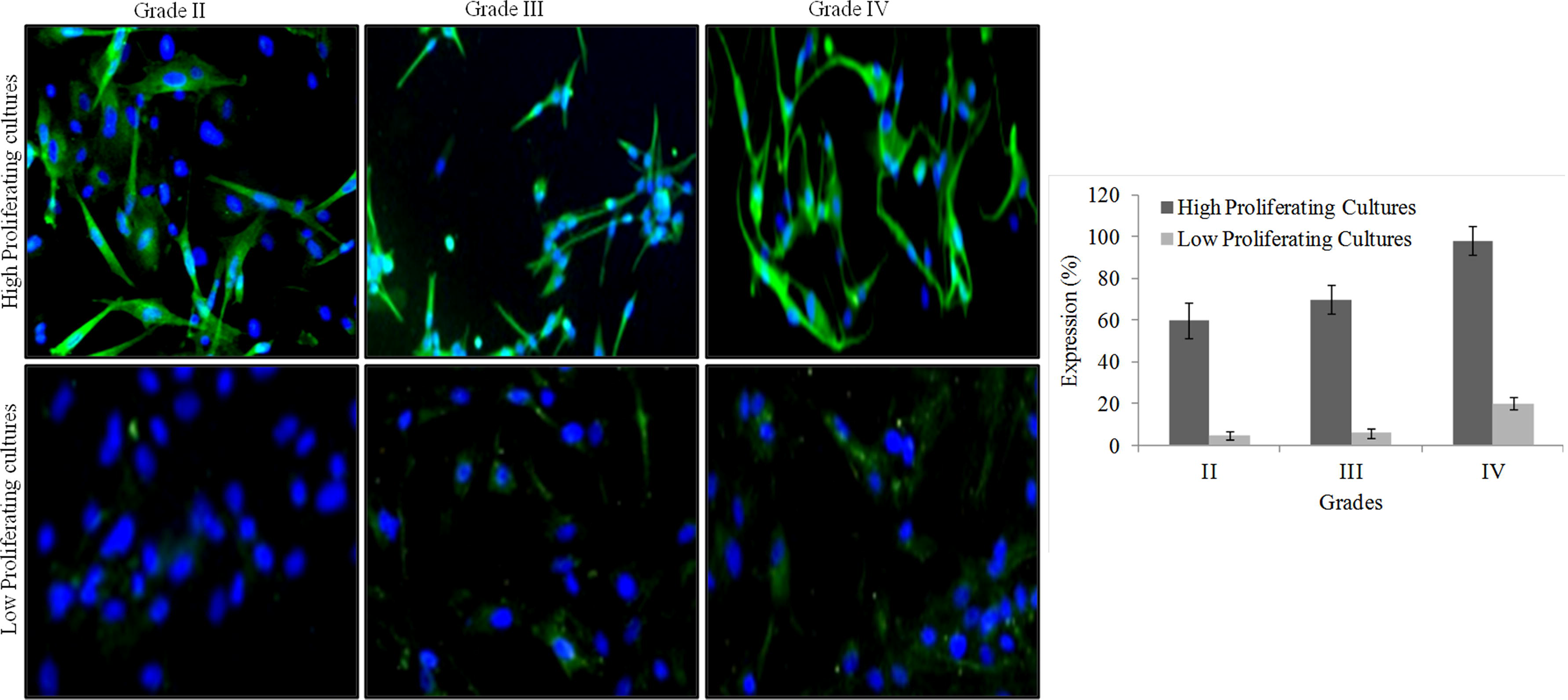
Representative Immuno-fluorescence staining of N-Cadherin. Fluorescent microscope images show the cell nuclei labeled with DAPI (Blue) and N-Cadherin labeled with respective antibody (Green, Alexaflour 488) and grouped grade-wise as per their proliferation rate. All images were captured at the Final Magnification of 100x.

**Figure 7:**
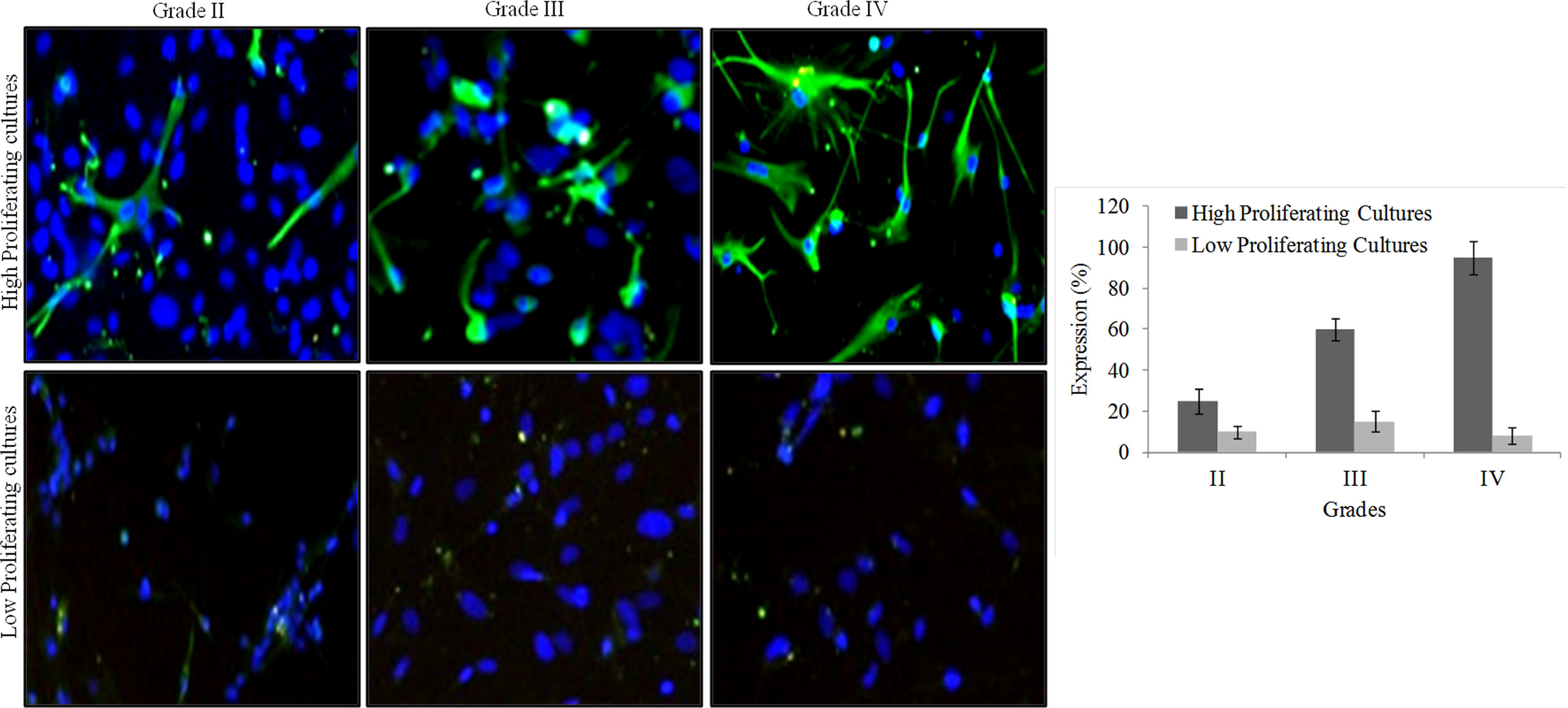
Representative Immuno-fluorescence staining of E-Cadherin. Fluorescent microscope images show the cell nuclei labeled with DAPI (Blue) and E-Cadherin labeled with respective antibody (Green, Alexaflour 488) and grouped grade-wise as per their proliferation rate. All images were captured at the Final Magnification of 100x.

**Figure 8:**
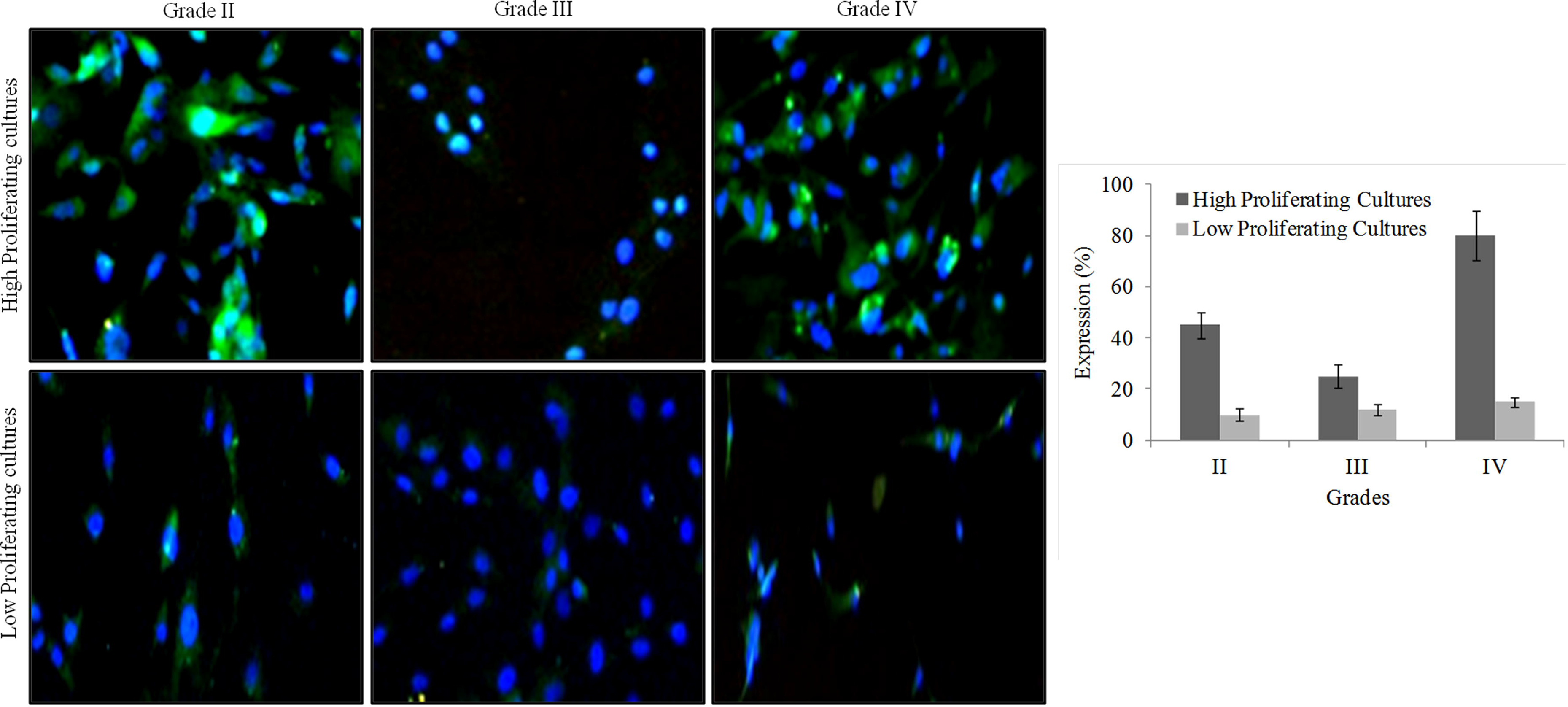
Representative Immuno-fluorescence staining of cMyc. Fluorescent microscope images show the cell nuclei labeled with DAPI (Blue) and cMyc labeled with respective antibody (Green, Alexaflour 488) and grouped grade-wise as per their proliferation rate. All images were captured at the Final Magnification of 100x.

### IDH1/2 mutation and 1p/19q co-deletion

IDH1/2 mutation and 1p/19q co-deletion status were evaluated in all samples, and their frequency concerning high and low-proliferating culture was determined (Supplementary Table 2). IDH1R132 mutation was detected in all samples from patients with grade II. Conversely, all grade IV samples were tested negative for IDH1/2 mutation, and only one sample from grade III showed positivity for IDH1R100. Complete 1p/19q co-deletion was detected in 6 of grade II, one of grade III, and none in grade IV samples.

### MIB1 and p53 expression in FFPE tissue section

MIB1 expression was found to be negative in all the grade II sample irrespective of proliferation status. However, all grade III and IV samples showed a differential degree of expression that were categorized as weak (1+), moderate (2+), and strong (3+), illustrated in Supplementary Table 1. In case of p53, proliferation status of culture appeared to play a considerable role as the number of positivity was more for those patients who had high-proliferating glioma cells in cultures (14 positives out of 19 samples) and less for those with low-proliferating cultures (6 positives out of 21 samples).

### Association of clinical and pathology features with the proliferation status of primary culture

Two-sided χ2 test to evaluate the association of proliferative ability of primary culture with age, sex, IDH mutation, 1p/19q co-deletion, p53, and MIB1 expression revealed that proliferation ability of primary culture was significantly associated with 1p/19q co-deletion (*p* = 0.0497) and p53 expression (*p* = 0.0044), but not with age, gender, IDH1/2 mutation and MIB1 expression (Table 2). Besides, simple linear regression analysis (Supplementary Figure 1) revealed that significant correlation was found in case of 1p/19q co-deletion (*p* = 0.05) and p53 expression (*p* = 0.05).

## Discussion

Therapeutic strategy and ensuing prognosis of gliomas differ considerably depending on the tumor grade and molecular biomarkers (15). Despite newer treatment modalities, the prognosis of glioma is poor, and survival statistics are not remarkable. The reason for treatment failure could be quiescent cells in tumor tissues that could later transform into a more aggressive phenotype, setting off tumor relapse (16). Unfortunately, the trigger for these transformations is poorly understood, and there is no therapeutic approach to delay or stop this process. Of late, *in vitro*, patient-specific glioma cultures have emerged as a standard model to get insight into these crucial aspects.

Although many studies pertaining to *in vitro* glioblastoma model have been published (11, 12, 17, 18), low-grade glioma models remain relatively unexplored. In this study, we successfully established *in vitro* primary culture of both low-grade and high-grade gliomas and evaluated their growth kinetics along with TMZ sensitivity and GMT expression pattern. Morphologically, there were no distinctive features noticed between low-grade and high-grade as both cultures showed a mixed cell population, mostly of dendritic and fibroblastic phenotypes. Intriguingly, there were conspicuous differences in the proliferation rate of glioma cultures, some being doubled swiftly in less than 72 h and others proliferating at a much lesser pace. We noticed this unique growth trend irrespective of tumor grades and subsequently categorized as high-proliferating and low-proliferating cultures. Moiseeva et al. (18) have reported the trend similar to our findings, albeit in the proliferation rate of high-grade glioma. In contrast, Mullins et al. (11) have reported no precise differences in the growth kinetics of cell lines established from fresh and frozen surgical materials from high-grade glioma.

It is well known that every tumor follows a well-defined logistic growth curve and differs in line with quiescent tumor cells (19). There was clear evidence of shortened doubling time, regardless of tumor grades, in a particular proportion of our studied patients. We presume that quiescent tumor cells that are not engaged in a cell cycle due to lack of oxygen or nutrients could have triggered into proliferation under *in vitro* culture conditions, which could plausibly indicate the impending progression of histological characteristics to a more aggressive phenotype.

Temozolomide (TMZ) is part of the standard of care for adjuvant therapy of malignant gliomas, despite the fact that at least 50% of the TMZ-treated patients eventually develop resistance to TMZ (20). Under such milieu, we intended to ascertain whether culture’s proliferative status could impact TMZ sensitivity. Thenceforth, we evaluated the IC50 of TMZ on all established glioma cultures. Low-proliferating cultures of grade II and IV tend to be more sensitive to TMZ with IC50 values lying within the tested concentration when compared with high proliferating cultures. However, all the grade III samples showed intense resistance to TMZ irrespective of their proliferation status. We presumed that high-proliferating cultures could have had drug-tolerant cells in a specific cellular state, allowing them to endure drug treatment and eventually giving rise to highly aggressive phenotype.

Mesenchymal differentiation or transformation contributes to a cellular ability to evade chemotherapies’ effects and promote drug resistance in several cancers (21, 22). Although the contribution of EMT in non-epithelial tumors like glioma is still debatable, we assume that GMT as a complement of EMT could be an essential process in glioma progression and promote drug resistance. Several studies have reported the modulation of distinctive EMT markers during the development of drug resistance in glioblastoma (7, 23). We also observed a notable variation in the expression profile of GMT markers amongst low-proliferating and high-proliferating cultures. Distinctive features among them were the strong expression of GFAP, N-cadherin, and E-cadherin in high-proliferating cultures regardless of tumor grade. While it has been suggested that the downregulation of E-cadherin and upregulation of N-cadherin is the hallmark of EMT (24), loss of E-cadherin may not be a necessary phenomenon for EMT as restoration of E-cadherin expression in E-cadherin negative malignant cells did not reverse the EMT process (25). Our findings conjecture that distinct GMT markers expression in high-proliferating culture could imply the invasive and aggressive nature of the tumor and related to TMZ resistance observed herein.

Another striking observation in our study was the nuclear expression of c-Myc in high-proliferating cultures as opposed to cytoplasmic expression in low-proliferating cultures. Although high expression of c-Myc was previously described in a large proportion of glioblastomas (26), information on cellular localization was sparse. Gurel et al. (27) have reported that increased nuclear expression of c-Myc could be a significant oncogenic event driving initiation and progression in human prostate cancer. Given its role in tumor progression, we speculate that nuclear cMyc expression could have a prognostic significance in gliomas.

Subsequently, we sought to assess the association of IDH1/2 mutation, 1p/19q co-deletion status, and other pathological features concerning the proliferation rate of cultured gliomas. We found a statistically significant association of 1p/19q co-deletion and p53 expression with proliferation rate. Since 1p/19q co-deletion has been demonstrated to have a significant protective effect on oligodendrogliomas (28), we presume that 1p/19q co-deletion could be associated with the proliferation status of primary glioma culture and have broad applicability in the prognosis of gliomas.

## Conclusion

In this study, we effectively demonstrated the practicality of the development of short-term glioma culture toward a functional approach for personalized medicine. Our study revealed the presence of highly proliferative, drug-resistant phenotypes with distinctive GMT phenotype irrespective of tumor grade. However, the clinical significance of differential growth kinetics, drug response, and distinct GMT phenotype observed in our study population need to be ascertained with active patients’ follow-up and survival analysis, which is underway in our laboratory. Nevertheless, our short-term glioma model could be a valuable tool for the management of low-grade and high-grade glioma and also provide cues about drug response and imminent tumor relapse.

## Ethics approval

This study was approved by the Institutional Research Advisory Board and Ethics Committee of KIMS Hospitals, and all procedures were performed following the Declaration of Helsinki, after obtaining written informed consent from the patients/legally accepted representatives.

## Consent to participate

Written informed consent was obtained from the patients/legally accepted representatives to participate in this study

## Consent for publication

All the patients who participated in this study has given consent for publication

## Availability of data and material

Not applicable

## Code availability

Not applicable

## Authors’ contributions

SSB and VKV designed the study. MKP preformed diagnosis, treatment, and management of patients. RK and AB collected the samples. SSB, VKV, and AB carried out experiments and acquired data. SM, RC, and SGR performed pathological and biochemical analysis. SSB and VKV performed analyses and interpretation of data. SSB and VKV wrote the manuscript, edited, and reviewed.

## References

1. Ostrom QT, Bauchet L, Davis FG, Deltour I, Fisher JL, Langer CE, et al (2014) The epidemiology of glioma in adults: a “state of the science” review. Neuro Oncol 16: 896–913

2. Yeole BB (2008) Trends in the brain cancer incidence in India. Asian Pac J Cancer Prev 9: 267–270

3. Louis DN, Perry A, Reifenberger G, Deimling A, Figarella-Branger D, Cavenee WK, et al (2016) The 2016 World Health Organization Classification of Tumours of the Central Nervous System: a summary. Acta Neuropathol 131:803

4. Diksin M, Smith SJ, Rahman R (2017) The Molecular and Phenotypic Basis of the Glioma Invasive Perivascular Niche. Int J Mol Sci 18: 2342

5. Behnan J, Ginocchiaro G, Hanna G (2019) The landscape of the mesenchymal signature in brain tumors. Brain 142: 847–866

6. Nowicki MO, Hayes JL, Chiocca EA, Lawler SE (2019) Proteomic Analysis Implicates Vimentin in Glioblastoma Cell Migration. Cancers 11: 466

7. Liang H, Chen G, Li J, Yang F (2019) Snail expression contributes to temozolomide resistance in glioblastoma. Am J Transl Res 11: 4277–4289

8. Elias MC, Tozer KR, Silber JR, Mikheeva S, Deng M, Morrison RS et al (2005) TWIST is expressed in human gliomas and promotes invasion. Neoplasia 7: 824–837

9. Noh MG, Oh SJ, Ahn EJ, Kim YJ, Jung TY, Jung S et al (2017) Prognostic significance of E-cadherin and N-cadherin expression in Gliomas. BMC Cancer 17: 583

10. da Hora CC, Schweiger MW, Wurdinger T, Tannous BA (2019) Patient-Derived Glioma Models: From Patients to Dish to Animals. Cells 8: 1177

11. Mullin CS, Schneider B, Stockhammer F, Krohn M, Classen CF et al (2013) Establishment and characterization of primary glioblastoma cell lines from fresh and frozen material: A detailed comparison. PLoS ONE 8: e71070

12. Ledur PF, Onzi GR, Zong H, Lenz G (2017) Culture conditions defining glioblastoma cells behavior: what is the impact for novel discoveries? Oncotarget 8: 69185–69197

13. Claus EB, Walsh KM, Wiencke JK, Molinaro AM, Wiemels JL, Schildkraut JM, et al (2015) Survival and low-grade glioma: the emergence of genetic information. Neurosurg Focus 38: E6

14. Greenwood SK, Hill RB, Sun JT, Armstrong MJ, Johnson TE, Gara JP, et al (2004) Population doubling: A simple and more accurate estimation of cell growth suppression in the in vitro assay for chromosomal aberrations that reduces irrelevant positive results. Environ Mol Mutagen 43:36–44

15. Olar A, Aldape KD (2012) Biomarkers classification and therapeutic decision-making for malignant gliomas. Curr Treat Options Oncol 13:417–436

16. Whittle IR (2004) The dilemma of low grade glioma. J Neurol Neurosurg Psychiatry 75 (Suppl II): ii31–ii36

17. Mullins CS, Schneider B, Lehmann A, Stockhammer F, Mann S, Classen CF, et al (2014) A Comprehensive Approach to Patient-individual Glioblastoma Multiforme Model Establishment. J Cancer Sci Ther 6: 177–187

18. Moiseeva NI, Susova OY, Mitrofanov AA, Panteleev DY, Pavlova GV, Pustogarov NA, et al (2016) Biochemistry (Moscow) 81: 628–635

19. Maurice Tubiana (1989) Tumor Cell Proliferation Kinetics and Tumor Growth Rate. Acta Oncologica 28:113–121

20. Lee SY (2016) Temozolomide resistance in glioblastoma multiforme. Genes Dis 3:198L210.

21. Prieto-Vila M, Takahashi RU, Usuba W, Kohama I, Ochiya T (2017) Drug Resistance Driven by Cancer Stem Cells and Their Niche. Int J Mol Sci 18: 2574

22. Scheel C, Eaton EN, Li SH, Chaffer CL, Reinhardt F, Kah KJ, et al (2011) Paracrine and autocrine signals induce and maintain mesenchymal and stem cell states in the breast. Cell 145:926–940.

23. Chen C, Han G, Li Y, Yue Z, Wang L, Liu J (2019) FOX01 associated with sensitivity to chemotherapy drugs and glial-mesenchymal transition in glioma. J Cellular Biochem 120: 882–893

24. Loh CY, Chai JY, Tang TF, Wong WF, Sethi G, Shanmugam MK, Chong PP, Looi CY (2019) The E-Cadherin and N-Cadherin Switch in Epithelial-to-Mesenchymal Transition: Signaling, Therapeutic Implications, and Challenges. Cells 8: 1118

25. Hollestelle A, Peeters JK, Smid M, Timmermans M, Verhoog LC, Westenend PJ, et al (2013) Loss of E-cadherin is not a necessity for epithelial to mesenchymal transition in human breast cancer. Breast Cancer Res Treat 138:47–57

26. Faria MH, Gonçalves BP, do Patrocínio RM, de Moraes-Filho MO, Rabenhorst SHB (2006) Expression of Ki-67, topoisomerase II alpha and c-MYC in astrocytic tumors: correlation with the histopathological grade and proliferative status. Neuropathol 26:519–527

27. Gurel B, Iwata T, Koh CM, Jenkins RB, Lan F, Dang CV, et al (2008) Nuclear MYC protein overexpression is an early alteration in human prostate carcinogenesis. Mod Pathol 21: 1156–1167

28. Hu N, Richards R, Jensen R (2016) Role of chromosomal 1p/19q co-deletion on the prognosis of oligodendrogliomas: A systematic review and meta-analysis. Interdisc Neurosurg 5: 58–63

