## Supplementary figures and images for "Patients’ Derived Short-Term Glioma Culture: Identification of Aggressive, Drug-Resistant Phenotype"

### Supplementary Figure 1

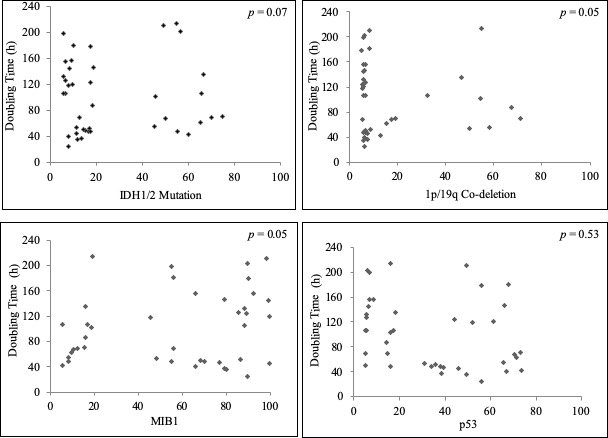
