## Supplementary Tables for "Patients’ Derived Short-Term Glioma Culture: Identification of Aggressive, Drug-Resistant Phenotype"

**Supplementary Table 1**

Clinical and Pathological Characteristics of the Glioma Patients

| **S. No** | **Age** | **Sex** | **Histology** | **Location** | **Grade** | **IDH1R132H** | **IDH1R132C** | **IDH1R100** | **IDH2R172** | **1p/19q**  **Co-deletion** | **P53** | **MIB1** |
| --- | --- | --- | --- | --- | --- | --- | --- | --- | --- | --- | --- | --- |
| 1. | 55 | M | Glioblastoma | Left temporal | IV | Negative | Negative | Negative | Negative | Negative | Negative | Positive (3+) |
| 2. | 35 | M | Diffuse astrocytoma | Left frontotemporal | II | Positive | Negative | Negative | Negative | Negative | Positive | Negative |
| 3. | 30 | F | Anaplastic astrocytoma | Right temporal | III | Negative | Negative | Negative | Negative | Negative | Positive | Positive (2+) |
| 4. | 29 | M | Oligodendroglioma | Right frontal | II | Positive | Negative | Negative | Negative | Positive | Negative | Negative |
| 5. | 39 | M | Diffuse astrocytoma | Right temporal | II | Positive | Negative | Negative | Negative | Positive | Positive | Negative |
| 6. | 55 | M | Glioblastoma | Right parietal | IV | Negative | Negative | Negative | Negative | Negative | Negative | Negative |
| 7. | 66 | F | Glioblastoma | Right parieto occipital | IV | Negative | Negative | Negative | Negative | Negative | Positive | Positive (2+) |
| 8. | 59 | M | Glioblastoma | Left fronto temporal | IV | Negative | Negative | Negative | Negative | Negative | Positive | Positive (3+) |
| 9. | 45 | M | Diffuse astrocytoma | Left frontal | II | Positive | Negative | Negative | Negative | Negative | Negative | Negative |
| 10. | 48 | F | Glioblastoma | Left temporal | IV | Negative | Negative | Negative | Negative | Negative | Negative | Positive (1+) |
| 11. | 19 | F | Anaplastic astrocytoma | Right parietal | III | Negative | Negative | Negative | Negative | Negative | Positive | Positive (2+) |
| 12. | 59 | M | Glioblastoma | Corpus callosal | IV | Negative | Negative | Negative | Negative | Negative | Negative | Positive (1+) |
| 13. | 63 | M | Glioblastoma | Right temporal | IV | Negative | Negative | Negative | Negative | Negative | Negative | Positive (2+) |
| 14. | 50 | M | Anaplastic astrocytoma | Right parietal | III | Negative | Negative | Negative | Negative | Negative | Positive | Positive (3+) |
| 15. | 71 | F | Glioblastoma | Right temporal | IV | Negative | Negative | Negative | Negative | Negative | Negative | Positive (2+) |
| 16. | 47 | F | Glioblastoma | Left parietal | IV | Negative | Negative | Negative | Negative | Negative | Negative | Positive (2+) |
| 17. | 37 | M | Diffuse astrocytoma | Right parieto occipital | II | Positive | Negative | Negative | Negative | Negative | Positive | Negative |
| 18. | 44 | M | Diffuse astrocytoma | Right temporoparietal | II | Positive | Negative | Negative | Negative | Positive | Positive | Negative |
| 19. | 38 | M | Oligo astrocytoma | Right frontoparietal | II | Positive | Negative | Negative | Negative | Positive | Negative | Negative |
| 20. | 69 | M | Glioblastoma | Right temporal | IV | Negative | Negative | Negative | Negative | Negative | Positive | Positive (1+) |
| 21. | 54 | M | Anaplastic astrocytoma | Right frontal | III | Negative | Negative | Positive | Negative | Positive | Positive | Positive (1+) |
| 22. | 38 | M | Diffuse astrocytoma | Right temporoccipital | II | Positive | Negative | Negative | Negative | Negative | Negative | Negative |
| 23. | 71 | M | Anaplastic astrocytoma | Left temporoparietal | III | Negative | Negative | Negative | Negative | Negative | Positive | Positive (1+) |
| 24. | 30 | M | Anaplastic astrocytoma | Right caudate | III | Negative | Negative | Negative | Negative | Negative | Positive | Positive (2+) |
| 25. | 53 | M | Glioblastoma | Left frontal | IV | Negative | Negative | Negative | Negative | Negative | Negative | Positive (2+) |
| 26. | 45 | M | Anaplastic astrocytoma | Right frontal | III | Negative | Negative | Negative | Negative | Negative | Positive | Positive (2+) |
| 27. | 63 | M | Glioblastoma | Left temporal | IV | Negative | Negative | Negative | Negative | Negative | Positive | Positive (3+) |
| 28. | 64 | M | Glioblastoma | Corpus callosum | IV | Negative | Negative | Negative | Negative | Negative | Negative | Positive (2+) |
| 29. | 47 | M | Oligodendroglioma | Left parietal | II | Positive | Negative | Negative | Negative | Positive | Negative | Negative |
| 30. | 27 | F | Diffuse astrocytoma | Right frontal | II | Negative | Positive | Negative | Negative | Negative | Negative | Negative |
| 31. | 57 | M | Glioblastoma | Left insular | IV | Negative | Negative | Negative | Negative | Negative | Positive | Positive (1+) |
| 32. | 79 | M | Glioblastoma | Right temporal | IV | Negative | Negative | Negative | Negative | Negative | Negative | Positive (1+) |
| 33. | 30 | F | Diffuse astrocytoma | Left frontal | II | Negative | Positive | Negative | Negative | Negative | Positive | Negative |
| 34. | 58 | F | Glioblastoma | Right temporal | IV | Negative | Negative | Negative | Negative | Negative | Negative | Positive (2+) |
| 35. | 37 | F | Oligodendroglioma | Right frontal | II | Positive | Negative | Negative | Negative | Positive | Negative | Negative |
| 36. | 67 | F | Glioblastoma | Left motor cortex | IV | Negative | Negative | Negative | Negative | Negative | Positive | Positive (1+) |
| 37. | 66 | F | Glioblastoma | Right parieto occipital | IV | Negative | Negative | Negative | Negative | Negative | Positive | Positive (2+) |
| 38. | 74 | M | Glioblastoma | Left fronto parietal | IV | Negative | Negative | Negative | Negative | Negative | Negative | Positive (1+) |
| 39. | 27 | M | Glioblastoma | Right temporal | IV | Negative | Negative | Negative | Negative | Negative | Positive | Positive (1+) |
| 40. | 56 | M | Glioblastoma | Left frontal | IV | Negative | Negative | Negative | Negative | Negative | Positive | Positive (2+) |
| 41. | 53 | F | Oligodendroglioma | Right Parietal | II | Positive | Negative | Negative | Negative | Positive | Negative | Negative |
| 42. | 43 | M | Anaplastic astrocytoma | Left frontal | III | Negative | Negative | Negative | Negative | Negative | Negative | Positive (2+) |
| 43. | 37 | M | Oligodendroglioma | Right temporoparietal | II | Positive | Negative | Negative | Negative | Negative | Positive | Negative |
| 44. | 54 | M | Glioblastoma | Right frontal | IV | Negative | Negative | Negative | Negative | Negative | Negative | Positive (3+) |
| 45. | 61 | M | Anaplastic astrocytoma | Right temporal | III | Negative | Negative | Negative | Negative | Negative | Positive | Positive (2+) |

Relevant clinical data concerning age (at the time of admission), gender (M = male; F = female), tumor histology, grade and localization, IDH1/2 mutation and 1p/19q co-deletion status along with presence/absence of proliferation marker MIB1 and Tumor Suppressor Marker p53 are summarized.

Supplementary Table 2

Frequency of IDH1/2 Mutations and 1p/19q Co-deletion Status in Malignant Glioma

| Mutations | Grade II | | Grade III | | Grade IV | |
| --- | --- | --- | --- | --- | --- | --- |
|  | HP Cultures | LP Cultures | HP Cultures | LP Cultures | HP Cultures | LP Cultures |
| Total Patients  IDH1R132H  IDH1R132C  IDH1R100  1p/19q codeletion | 7  6  1  0  2 | 5  4  1  0  4 | 4  0  0  1  1 | 3  0  0  0  0 | 8  0  0  0  0 | 13  0  0  0  0 |

HP Cultures – High Proliferating Cultures; LP Cultures – Low Proliferating Cultures

Supplementary Table 3

Association of Clinical and Molecular Pathology Features of Malignant Glioma with Proliferation rate of Primary Culture

|  | High Proliferation Culture | Low Proliferation Culture | χ2 | *p value* |
| --- | --- | --- | --- | --- |
| Total number of cases  Gender (Female/Male)  Age at diagnosis (years)  IDH1R132 (Positive/Negative)  IDH1R132H (Positive/Negative)  IDH1R132C (Positive/Negative)  IDH1R100 (Positive/Negative)  1p/19q co-deletion (Positive/Negative)  p53 (Positive/Negative)  MIB1 (Positive/Negative) | 19 (47%)  4/15  52.53 ± 15.45  7/12  6/13  1/18  1/18  2/17  14/5  12/7 | 21 (53%)  8/13  47.77 ± 15.17  3/18  4/17  1/20  0/21  6/15  6/15  15/6 | –  1.3796  –  2.7068  1.6708  0.0053  0.4699  3.9665  8.1203  0.3110 | –  0.2402  0.3316**^$^**  0.0999  0.1961  0.9421  0.4930  0.0497*  0.0044*  0.5770 |

**^$^** Students ‘t’ test; * statistically significant at *p* < 0.05
